# Clathrin recruitment and assembly of phosphorylated B cell receptor is impaired in human diffuse large B cell lymphoma

**DOI:** 10.64898/2026.09.22.753512

**Authors:** Aleah D. Roberts, Kem A. Sochacki, Louis M. Staudt, Justin W. Taraska

## Abstract

The B cell receptor (BCR) drives the differentiation of naive B cells into activated plasma cells that produce antibodies. The BCR consists of a plasma membrane-bound surface immunoglobulin (sIg) that binds antigens, and two coreceptor subunits CD79A and CD79B. The intracellular domains of CD79A and CD79B initiate immune signaling and endocytosis of the receptor in healthy B cells. In diffuse large B cell lymphoma (DLBCL), an aggressive form of human blood cancer, the activated B cell like (ABC) subtype exhibits constitutive signaling that drives survival and proliferation. Here, in a model of human ABC DLBCL, we determine the localization of BCR components relative to plasma membrane structures using correlative super-resolution light and platinum replica transmission electron microscopy. We find that spontaneous clusters of the surface Immunoglobulin common to ABC lymphoma localize to smooth raised membrane domains. These structures are involved in the endocytosis of large receptor clusters. Surprisingly, activated phosphorylated CD79A (pCD79A) shows little colocalization with clathrin on or off smooth raised membranes. Furthermore, pCD79A spatially segregates away from both the slg and CD79B subunits of the BCR complex. A similar distribution was found for downstream phosphorylated Src family kinases. We propose that a disengagement of pCD79A away from sIg and CD79B allows the receptor to evade down-regulation through endocytosis and maintains signaling at the plasma membrane. This mechanism could drive aberrant signaling and the proliferation of lymphomas leading to disease

## INTRODUCTION

B cell lymphocytes are key components of the immune system that produce antigen-specific antibodies to combat infections. When the immune system detects a foreign antigen, B cells engage extracellular antigens through the plasma membrane localized B cell receptor (BCR) (1, 2). The BCR acts as a multiprotein complex consisting of three subunits: the surface immunoglobulin (sIg) and two co-receptor proteins, CD79A and CD79B (3, 4) (Fig. 1A). During immune activation, antigen binding induces crosslinking of multiple BCRs into large macroscopic clusters at the plasma membrane. This clustering triggers phosphorylation of the intracellular domains of CD79A and CD79B by recruited Src family kinases. Src kinases then drive a cascade of phosphorylation events that initiate an inflammatory response and trigger endocytosis of the B cell receptor to terminate signaling (5, 6). In disease, chronically active inflammatory signaling from the BCR is observed in Diffuse Large B Cell Lymphoma (DLBCL), a fast-growing human cancer, and drives aggressive proliferation of neoplastic B cells (7–10). We hypothesized that dysregulation of endocytic mechanisms for BCRs and receptor organization that normally terminate inflammatory signaling may contribute to chronic activation in lymphoma.

**Figure 1.**
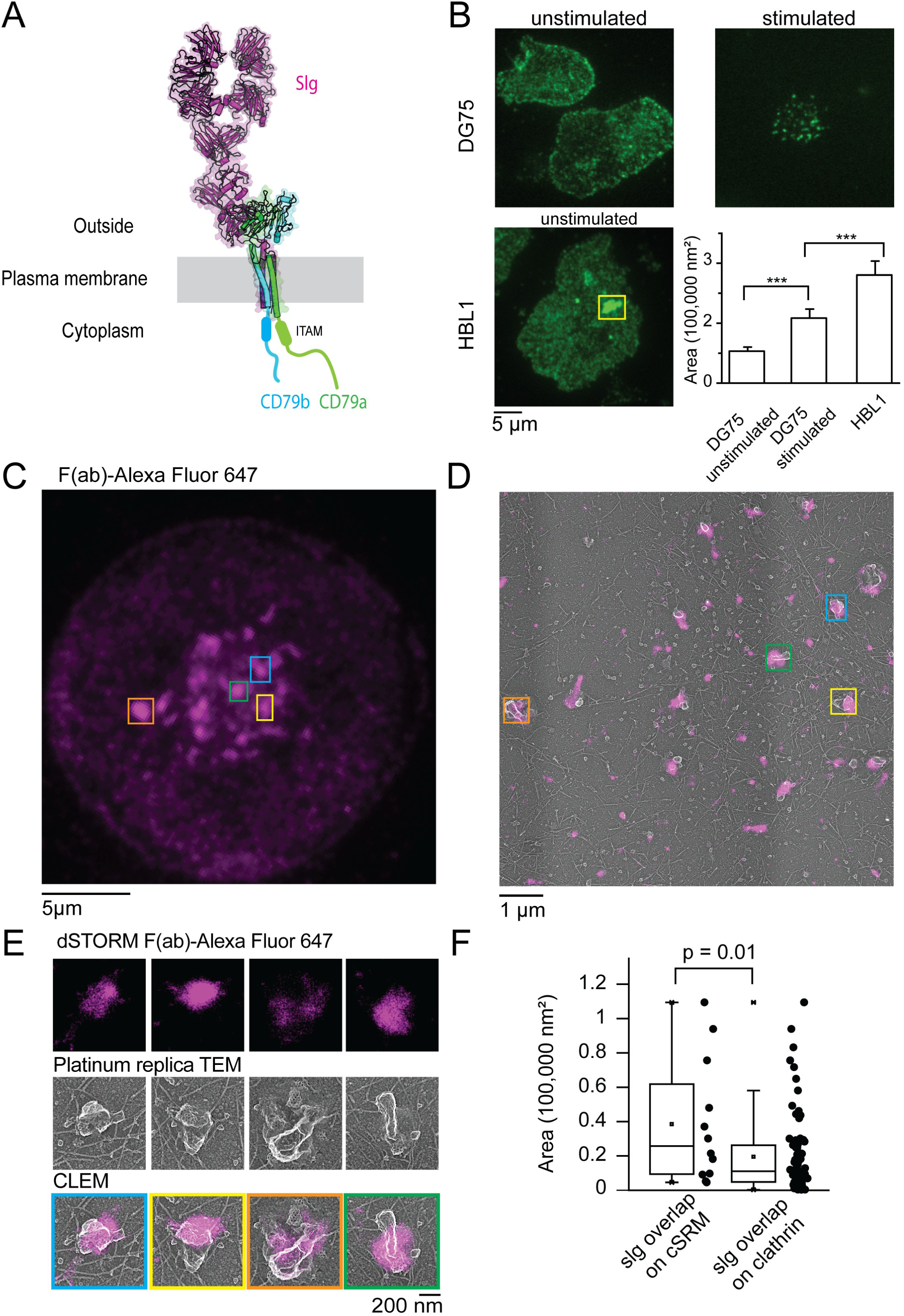
(A) Cartoon of the B cell receptor (BCR) at the plasma membrane (PDB: 7WSP). (B) TIRF images of surface immunoglobulin (sIg) in DG75 and HBL1 cells and quantification of sIg cluster area (n = 10 unstimulated DG75 cells, 11 stimulated DG75 cells, and 20 HBL1 cells). (C) TIRF image of an HBL1 cell highlighting regions containing prominent BCR clusters. (D) Correlated dSTORM and platinum replica TEM image of the indicated region. (E) Enlarged views of selected regions from D. (F) Quantification of BCR fluorescence associated with clathrin-containing membrane structures or clathrin alone (n = 7 cells from 3 biological replicates).

The BCR complex and its ligands in B cells are internalized from the plasma membrane by several distinct endocytic mechanisms to accommodate the diverse properties of antigenic stimuli (11). Specifically, antigens vary in size, concentration, avidity, or membrane binding properties that all present unique biophysical challenges for endocytosis. For example, antigens present on the surface of bacteria or complexed with an adjuvant in a vaccine are large and thought to be internalized by phagocytosis (12, 13). Membrane-bound antigens on the surface of an antigen presenting cell are internalized though trogocytosis (14, 15). Soluble antigens at low concentrations are taken up through clathrin-mediated endocytosis or fast endophilin-mediated endocytosis (FEME) (16–18). Finally, high concentrations of soluble antigen that induce large clusters of activated receptors are internalized by clathrin that resides on large smooth raised membrane structures (cSRM) through a hybrid form of endocytosis (19).

In this study, we used advanced microscopies to visualize BCR clustering and signaling in models of DLBCL to study the role of clathrin and cSRM mediated endocytosis in chronic BCR activation in human cancers. DLBCL is the most commonly-diagnosed non-Hodgkins lymphoma (NHL) in humans and accounts for ∼40% of NHL cases worldwide (20). There are two major subtypes—activated B cell like (ABC) and germinal center B cell like (GCB) (21, 22). These have been classically distinguished by gene expression profiles. ABC DLBCL express genetic characteristics of B cells that have been activated through BCR signaling (i.e., genes related to NF-κB), while the profile of GCB DLBCL cells is similar to inactive germinal center B cells. Classic work has shown that ABC DLBCL cells have clusters of BCR in the absence of foreign antigen and rely on stimulation by self-antigens for survival and proliferation (7, 23). Recent data showed that blocking BCR glycosylation— whether genetically or chemically— disperses BCR clusters in ABC DLBCL thus attenuating NF-κB activation (24). Together, these studies suggest that BCR clusters play a key role in the maintenance of pathogenic signaling. Yet, the structure and mechanism of how steady-state BCR clusters are maintained at the plasma membrane to alter signaling and endocytosis in this type of cancer are mostly unknown.

Here, using nanoscale correlative imaging, we find that clusters of the surface immunoglobulin (slg) in unstimulated ABC DLBCL cell line HBL1 localize to large clathrin-coated smooth, raised membrane structures (previously described cSRM structures, (19)). These large inward plasma membrane invaginations are involved in endocytosis of activated BCR clusters (19). In contrast, we observe that the phosphorylated (activated) CD79A subunit of the BCR can localize to cSRMs but shows minimal localization with clathrin, a pattern not observed in other B cell types. This defect in clathrin association could allow the activated subunit to avoid downregulation through endocytosis. Analysis of individual components of the BCR complex (sIg, CD79A, and CD79B; Fig. 1A) in HBL1 cells further indicates that while sIg, CD79A, and CD79B colocalize, the phosphorylated form of CD79A segregates away from the sIg/CD79B complex. Together, these results provide new insights into the stability, distribution, and trafficking of the activated BCR complex at the plasma membrane of blood cancers.

## RESULTS

### sIg clusters in HBL1 cells colocalize with clathrin on smooth raised membrane domains

Quiescent B cells do not exhibit strong BCR clustering at the plasma membrane and the receptor is mostly distributed evenly across the plasma membrane (Figure 1 B top row, left side) (2, 25–27). In some antigen stimulated cells, however, the BCR forms large clusters due to the crosslinking of the sIg domain (Figure 1 B top row, right side). B cell lines and B cell samples from patients with activated B cell (ABC) like DLBCL cancers, however, maintain large BCR clusters at the plasma membrane even without foreign antigen engagement (Figure 1 B bottom row, yellow box high lights a BCR cluster in the HBL1 cell line) (7, 23). These are sometimes called “spontaneous clusters” to distinguish them from clusters activated by a foreign extracellular antigen. Figure 1B shows that spontaneous slg clusters in HBL1 cells were significantly larger than both unstimulated or stimulated DG75 cells (Figure 1 B, bar graph).

In order to investigate endocytic proteins that colocalize with spontaneous BCR clusters and may contribute to reduced sIg uptake, we used super-resolution correlative light and platinum replica transmission electron microscopy to determine what type of plasma membrane endocytic structures underly sIg clusters in HBL1 cells (29). First, we stained cells with a monovalent F(ab) fragment to identify the sIg and then unroofed the cells to expose the inner plasma membrane. We then performed super-resolution dSTORM imaging of the sIg before processing samples for platinum replica transmission electron microscopy (TEM). Correlated fluorescence and TEM images showed that spontaneous sIg clusters are often localized with clathrin lattices that reside on distinctive smooth raised membrane domains (cSRMs, Fig. 1C - E). Smooth raised membrane domains are characterized by large inward plasma membrane invaginations that appear devoid of rough texture commonly found on the plasma membrane of these cells and that lack identifiable coat proteins (19). These structures are uniquely found in activated B cells and are thought to endocytose large BCR clusters. Figure 1D and E show correlated fluorescence and CLEM images of an HBL1 cell with sIg clusters highlighted in colored boxes. In these images, roughly 80% of the sIg clusters identified in TIRF correlated with cSRM structures (calculation described in methods). Figure 1E shows zoomed images of sIg clusters overlapping cSRMs. We next measured the size of the sIg clusters that overlap with cSRMs. To do this, we generated segmented maps of clathrin and smooth raised membrane structures and overlayed these regions with the reconstructed super-resolution fluorescence. We calculated the area of sIg clusters that touch either clathrin on cSRM structures or clathrin in other plasma membrane areas (Fig. 1F). SIg clusters that overlapped with cSRM structures were significantly larger (average area of 38,562 nm^2^ ± 10,407 nm^2^) than fluorescence associated with clathrin alone (average area of 19,484 nm^2^ ± 2,368 nm^2^).

These CLEM data confirm that 1) cSRM structures are present in HBL1 cells and, 2) that they are strongly correlated with particularly large sIg clusters (spontaneous clusters) that form at the plasma membrane of HBL1 cells. Because cSRMs are involved in the uptake of activated BCR clusters into the cytosol, we next tested if the these clusters contain active or inactive BCR receptors (19).

### Phosphorylated CD79A is not colocalized with clathrin

Immunoreceptor tyrosine activation motifs (ITAMs) on the cytoplasmic tails of CD79A and CD79B are phosphorylated by Src family kinases after antigen engagement of the sIg to activate immune signaling and recruit clathrin to the activated receptor (9). The activated (i.e. phosphorylated) form of CD79A can be detected with an antibody that binds to the phosphorylated tyrosine residue in the membrane proximal ITAM of CD79A. Thus, we correlated the sIg (green, diffraction limited TIRF), pCD79A (magenta, dSTORM), and plasma membrane structures (TEM images) to test if sIg clusters that colocalized with cSRMs also contain activated phosphorylated CD79A. Figure 2A shows a TIRF image of two HBL1 cells with prominent sIg clusters highlighted in colored boxes. We used TIRF images to identify prominent sIg clusters in the cell and then observed pCD79a localization and plasma membrane structures at the same location (Fig. 2B). Surprisingly, unlike slg, pCD79A is often localized in clusters on smooth raised membranes but these clusters do not regularly overlap with clathrin. Honeycomb clathrin lattices are colored in yellow transparency in Figure 2B.

**Figure 2.**
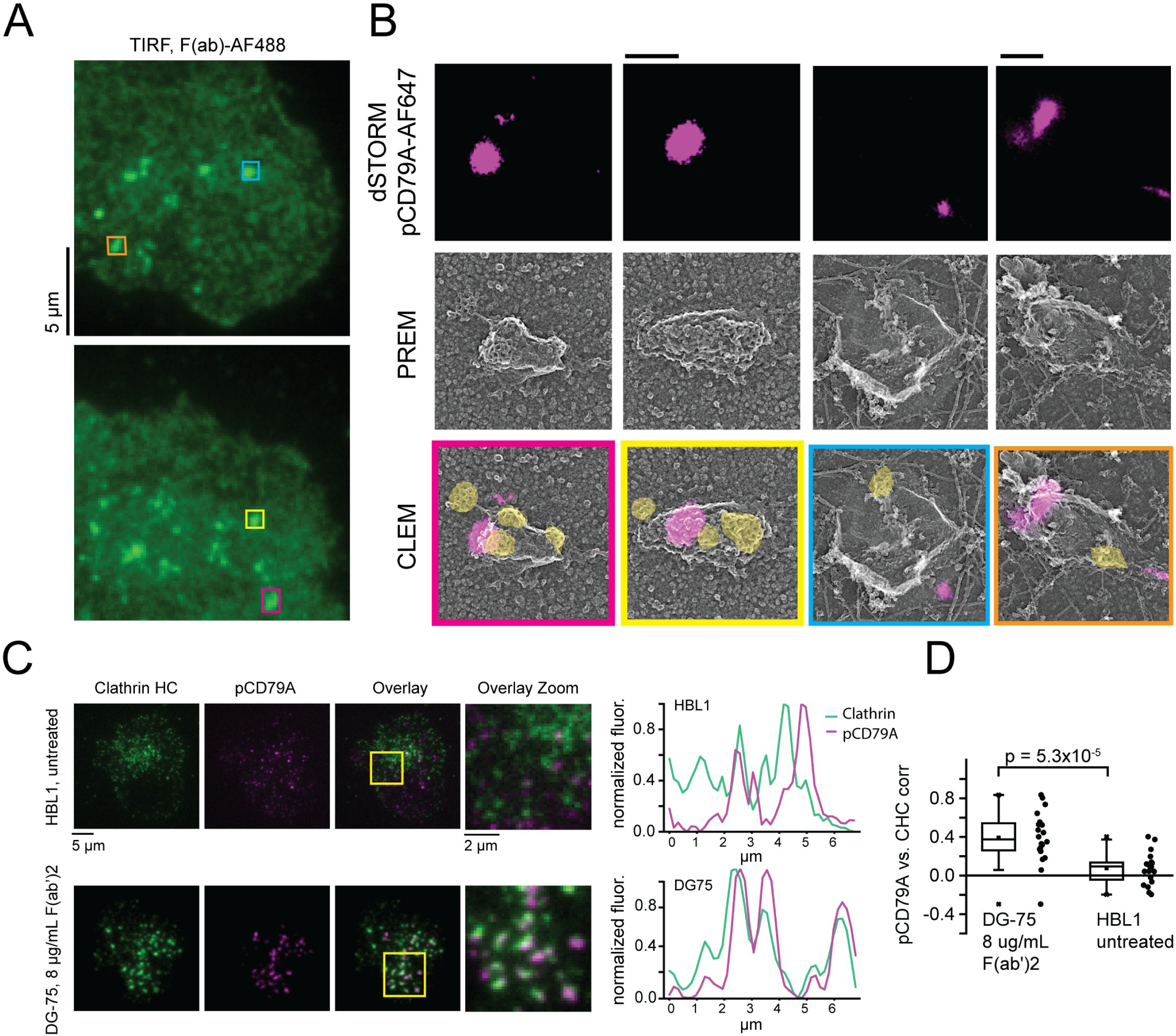
**(A)** TIRF images of HBL1 cells stained with a F(ab) fragment. Prominent clusters are highlighted in color coded boxes. (**B)** Zoomed images of clusters shows phosphorylated CD79A is present on cSRMs but does not spatially overlap with clathrin. Yellow transparency are segmented clathrin regions. Scale bars are 200 nm. **(C)** Colocalization analysis of pCD79A and Clathrin Heavy Chain in HBL1 cells and stimulated DG75 cells. (**D)** Colocalization values between pCD79A and CHC (n = 20 cells per sample, with 260 spots analyzed from HBL1 cells and 104 spots from DG75 cells) in untreated HBL1 and stimulated DG75. pCD79A were selected as the reference to determine the level of CHC colocalization.

To test whether clathrin recruitment to pCD79A is impaired across the plasma membrane, we imaged clathrin heavy chain (CHC) and pCD79A. This allowed us to collect data from a larger number of cells at a more wide-scale optical resolution compared to EM imaging. Figure 2C shows representative images of pCD79A and CHC staining in HBL1 cells along with control stimulated DG75 cells. As expected, after stimulation, clathrin is strongly recruited to the pCD79A protein in DG75 cells. In contrast, in HBL1 cells at rest, where CD79A is phosphorylated due to the chronic nature of activation, pCD79A does not substantially colocalize with clathrin. Figure 2D shows quantification of the colocalization analysis collected for all clathrin spots across many cells and replicates. We conclude that while pCD79A in DG75 cells colocalizes strongly with clathrin, HBL1 cells appear to be deficient in clathrin association with pCD79A specifically.

Several ABC DLBCL cell lines and patient samples express CD79B mutations that may affect endocytosis sIg clusters. For example, HBL1 cells express a heterozygous mutant: CD79B^Y196F^(7). The Y196F mutation alters the binding site for the clathrin adaptor AP-2 and similar mutations at this location have been shown to block endocytosis (28). Thus, we hypothesized that resting HBL1 cells may have a reduced rate of sIg recycling from the plasma membrane due to this or other mutations. To test for differences in receptor endocytosis, we measured uptake of the sIg in resting HBL1 cells. We observed HBL1 cells had lower levels of sIg uptake from the plasma membrane, compared to the DG75 control. Figure 3A shows data at the 10 and 20-minute time point, but the trend continues for up to an hour (Supp. Fig. 1). This reduced rate of sIg uptake may be due to reduced clathrin-mediated or clathrin-independent endocytosis.

**Figure 3.**
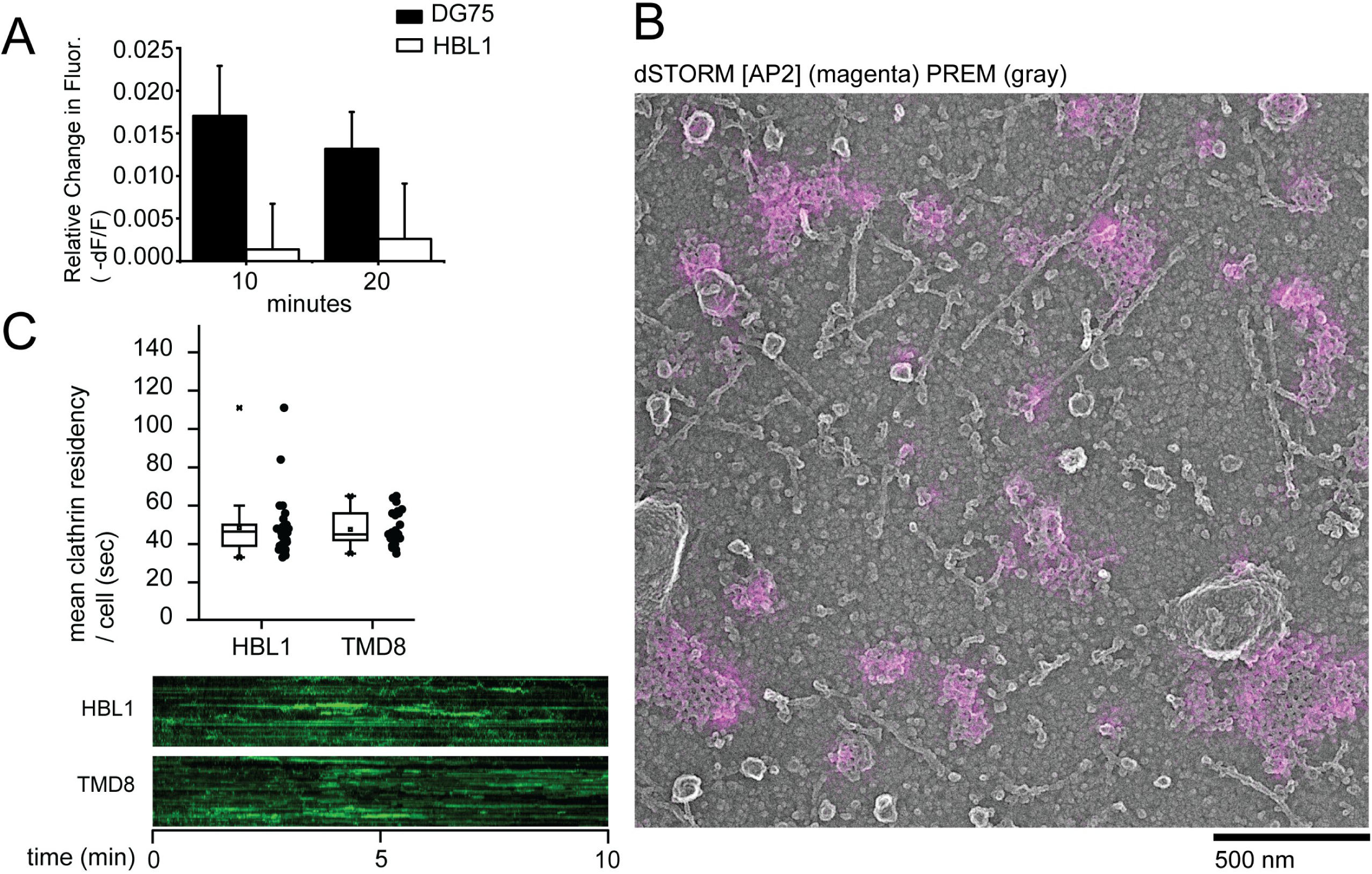
**(A)** Flow cytometry analysis of surface immunoglobulin (sIg) internalization over time in DG75 and HBL1 cells (n = 5 biological replicates for DG75 and 3 biological replicates for HBL1). **(B)**Correlated super-resolution dSTORM fluorescence of AP-2 (magenta) and platinum replica TEM of an HBL1 cell inner plasma membrane. **(C)** Clathrin residency time of HBL1 and TMD8 cells generated from live-cell TIRF imaging. An example kymograph for each cell type shows fluorescence projected through time for an example region. (n = 3 biological replicates, with 10 cells analyzed per experiment)

To test if defects in AP2 prevents clathrin recruitment to pCD79A, we imaged AP2 and clathrin colocalization at the plasma membrane using CLEM (Fig. 3B). AP2 is highly colocalized with clathrin in HBL1 at the nanoscale. Given that AP2 can recruit clathrin to the plasma membrane in HBL1 cells, the segregation of pCD79A away from clathrin is not a result of a mutation induced defect in AP2 function or clathrin assembly in HBL1 cells.

Next, we tested if defects in clathrin recruitment to pCD79A might be caused by defects in other aspects of clathrin mediated endocytosis. Specifically, we analyzed clathrin residency time in live cells to assess endocytic function in two ABC cell types (HBL1 and TMD8). HBL1 and TMD8 cells had clathrin residency times of around 50 seconds (Fig. 3C). These times are within the normal range across many standard cell types and suggest that clathrin mediated endocytosis is generally functioning within normal ranges in these cell lines and is not grossly impaired (30, 31).

Finally, to investigate whether cSRM structures play a role in maintaining sIg clusters at the plasma membrane, we treated HBL1 cells with the actin inhibitors Cytochalasin D or Latrunculin A to reduce the density of cSRMs at the plasma membrane as we previously reported in stimulated DG75 cells (19). We imaged sIg to determine the effect on receptor clustering. None of the actin inhibitor treatments we tested, however, reduced the density of cSRMs at the plasma membrane of HBL1 cells (Supp. Fig. 2). This could be because actin is required for formation of the cSRM structures but not their maintenance, and so treatment with actin inhibitors is most effective before receptor activation but is not effective in HBL1 cells, which are constitutively activated (19).

Together these data support the model that spontaneous clusters of sIg in HBL1 cells localize on plasma membrane structures that typically endocytose activated receptor clusters (cSRMs specifically). In HBL1 cells, however, the pCD79A does not colocalize strongly with clathrin. These data lead us to look more closely at the localization of plasma membrane associated pCD79A.

### Phosphorylated CD79A segregates away from the sIg and CD79B complex

In our correlative imaging studies, pCD79A does not substantially colocalize with the sIg. To further investigate this result, we compared the pattern of pCD79A fluorescence in HBL1, to pCD79A localization in DG75 cells as a control. As expected, without antigen, there is little to no visible staining for pCD79A in DG75 cells (Fig. 4A top panel). After antigen stimulation, however, a robust signal for pCD79A is highly colocalized with the sIg (Fig. 4A bottom panel). The lack of pCD79A staining in unstimulated DG75 cells also indicates that there are likely no off-target proteins stained by the pCD79A antibody. HBL1 cells are constitutively active and contain positive staining for pCD79A without stimulation. Therefore we expected unstimulated HBL1 cells to contain pCD79A staining that appears similar to stimulated DG75 cells. Figure 4B (and Supp. Fig. 3) shows regions in HBL1 that have peaks of pCD79A fluorescence with no substantial corresponding peak at the same location for the sIg (highlighted in yellow boxes in Fig. 4B, top panel). We also observed a similar localization pattern for pCD79A and sIg in TMD8 cells, another ABC DLBCL cell line (Fig. 4B bottom panel). The visible colocalization and measured correlation coefficients between sIg and pCD79A are substantially lower in TMD8 and HBL1 than stimulated DG75 cells (Fig. 4D). In addition to whole-cell correlation analysis shown in Figure 4D, we also analyzed the correlation between individual pCD79A peaks and the corresponding sIg fluorescence using a previously described high-throughput automatic method for analyzing pairwise correlation data (32). Here, we analyzed hundreds of pCD79A fluorescent spots across multiple cells and consistent with our whole-cell analysis found significantly less colocalization of pCD79A with sIg, compared to stimulated DG75 cells (Fig. 4E). The unique pCD79A localization observed in both HBL1 and TMD8 cells, suggests that pCD79A separation from the sIg may be a generalizable feature of ABC DLBCL lymphoma.

**Figure 4.**
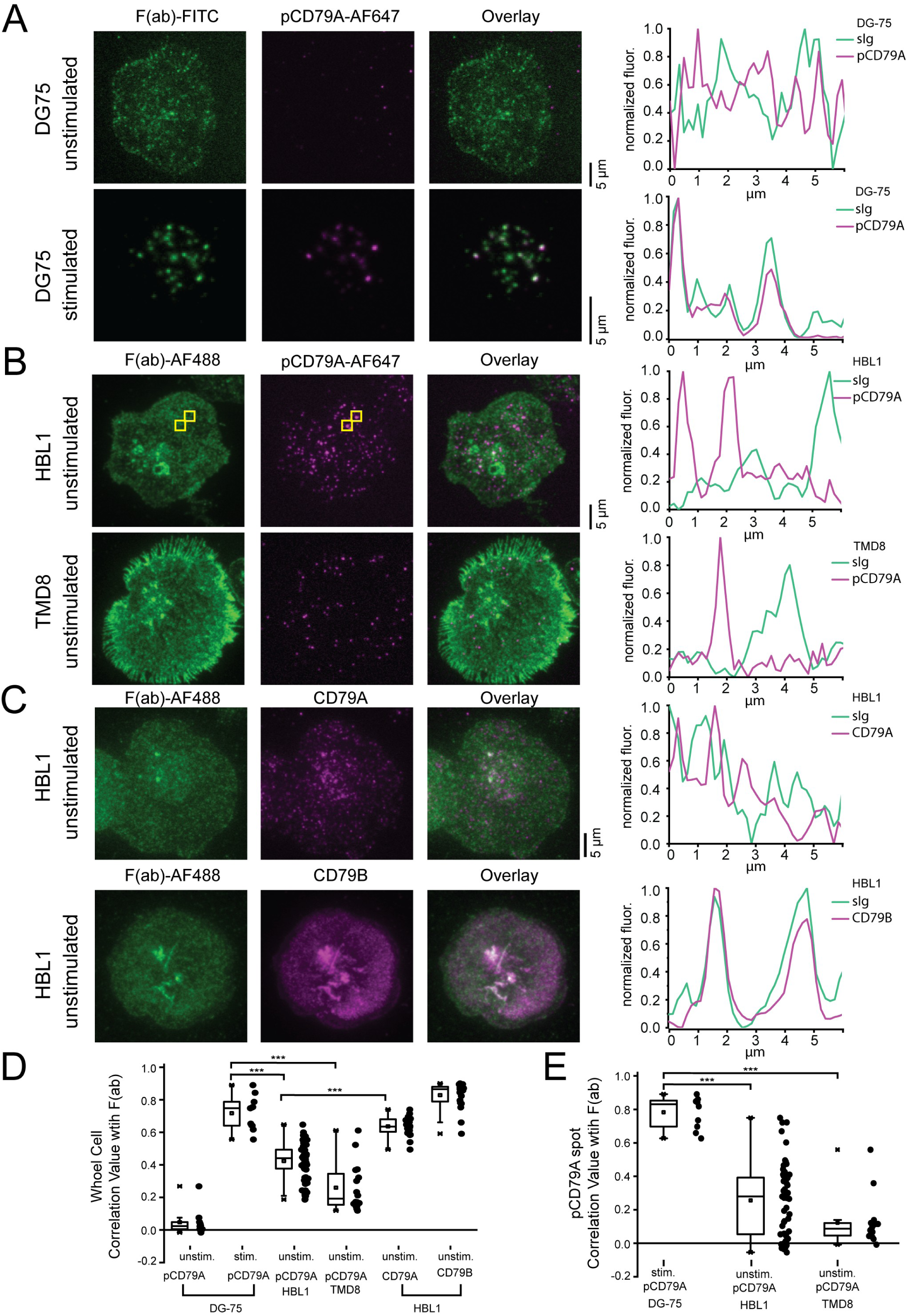
**(A)** TIRF images of sIg and pCD79A DG75 cells **(B)** TIRF images of sIg and pCD79A in HBL1 and TMD8 cells **(C)** TIRF images of CD79A and CD79B in HBL1 cells (**D)** Whole cell colocalization values of the sIg compared to staining for pCD79A, CD79A, or CD79B. (DG75, unstimulated n = 10 cells, DG75 stimulated n = 11 cells, both collected in 1 biological replicate; HBL1 unstimulated n = 61 cells collected over 3 biological replicates; TMD8 unstimulated n = 16 cells collected in 1 biological replicate; HBL1, unstimulated CD79A correlation n = 20 collected in 1 biological replicate; HBL1, unstimulated CD79B correlation n = 20 collected in 1 biological replicate) (**E)** Colocalization values for individual pCD79A spots correlated with sIg. pCD79A spots were selected as a reference to evaluate colocalization with sIg. (DG75 stimulated n = 116 spots analyzed; HBL1 unstimulated n = 4,078 spots analyzed; TMD8 unstimulated n = 821 spots analyzed)

To further explore the localization of pCD79A, we analyzed the spatial distributions of each component of the BCR complex. In HBL1 cells, CD79B was highly colocalized with the sIg (Fig. 4C bottom panel). The correlation values for CD79B and sIg were also high, suggesting that CD79B is complexed with sIg as expected in HBL1 cells. Total CD79A however, had lower colocalization with the sIg, compared to CD79B (Fig. 4C top panel, and correlation values in part D). This is consistent with the above data because the CD79A antibody binds both the inactive and phosphorylated CD79A protein, which is not colocalized. Thus, we would expect a lower overall correlation in these two probes. From these data we conclude that the non-phosphorylated CD79A protein is complexed with sIg/CD79B, while the activated phosphorylated CD79A protein separates away from sIg and CD79B in HBL1 cells. Colocalization between pCD79A and sIg is significantly less than colocalization between CD79A and sIg, suggesting that phosphorylation may be the trigger for dissociation of CD79A. To confirm these results, we further analyzed the localization of pCD79A in Figure 5 using super-resolution structured illumination microscopy (SIM). In unstimulated HBL1 cells, again we observed multiple cellular regions that contained clustered pCD79A (magenta) without significant colocalization of sIg (green) (Fig. 5A). In contrast, stimulated DG75 cells (Fig. 5B) have clustered pCD79A that is highly colocalized with sIg. Correlation analysis of SIM data found that pCD79a is significantly less correlated with the sIg in HBL1 compared to stimulated DG75 cells (Supp. Fig. 4). Interestingly, HBL1 cells do have a higher baseline level of diffuse and small clustered domains in comparison to the highly clustered receptors in active DG75 cells that have very little un-clustered signal. These data suggest that the slg complex in HBL1 cells could be more heterogeneously distributed than strongly activated BCRs in other B cell types.

**Figure 5.**
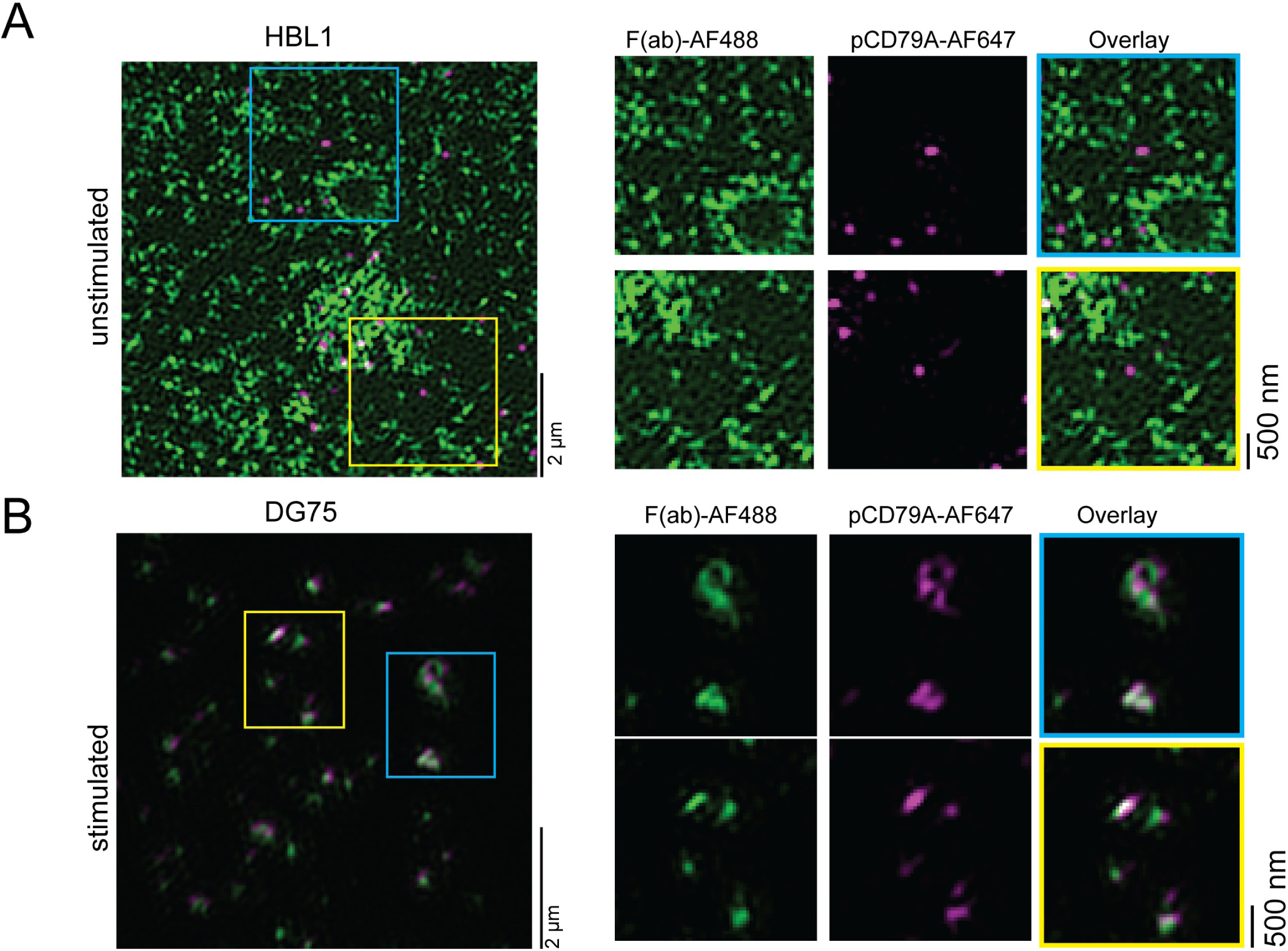
**(A)** TIRF SIM images of sIg (green) and pCD79a (magenta) in an unstimulated HBL1 cell. Colored boxes show zoomed in regions of the cell. (n = 25 cells, from 2 biological replicates) **(B)** TIRF-SIM image of sIg (green) and pCD79A (magenta) in a stimulated DG75 cell. Colored boxes show zoomed regions. (n = 43 cells, from 2 biological replicates)

Internalization of active BCR is regulated by both CD79A and CD79B (28), and segregation of the phosphorylated CD79A away from the BCR complex could contribute to the low levels of clathrin recruitment, providing a mechanism for the activated CD79A protein to evade down-regulation by endocytosis and maintain aberrant signaling. To determine whether the pCD79A protein can activate classic downstream signaling, we imaged with an antibody against activated phosphorylated (Y416) Src family kinase members (pSFK) (Figure 6). In control DG75 cells, stimulation induced pSFK that was strongly colocalized with sIg clusters. HBL1 cells, however, contain pSFK signal that was poorly colocalized with the sIg, similar to pCD79A localization. These data suggest that the disengaged pCD79A can still activate Src family kinases downstream of receptor activation.

**Figure 6.**
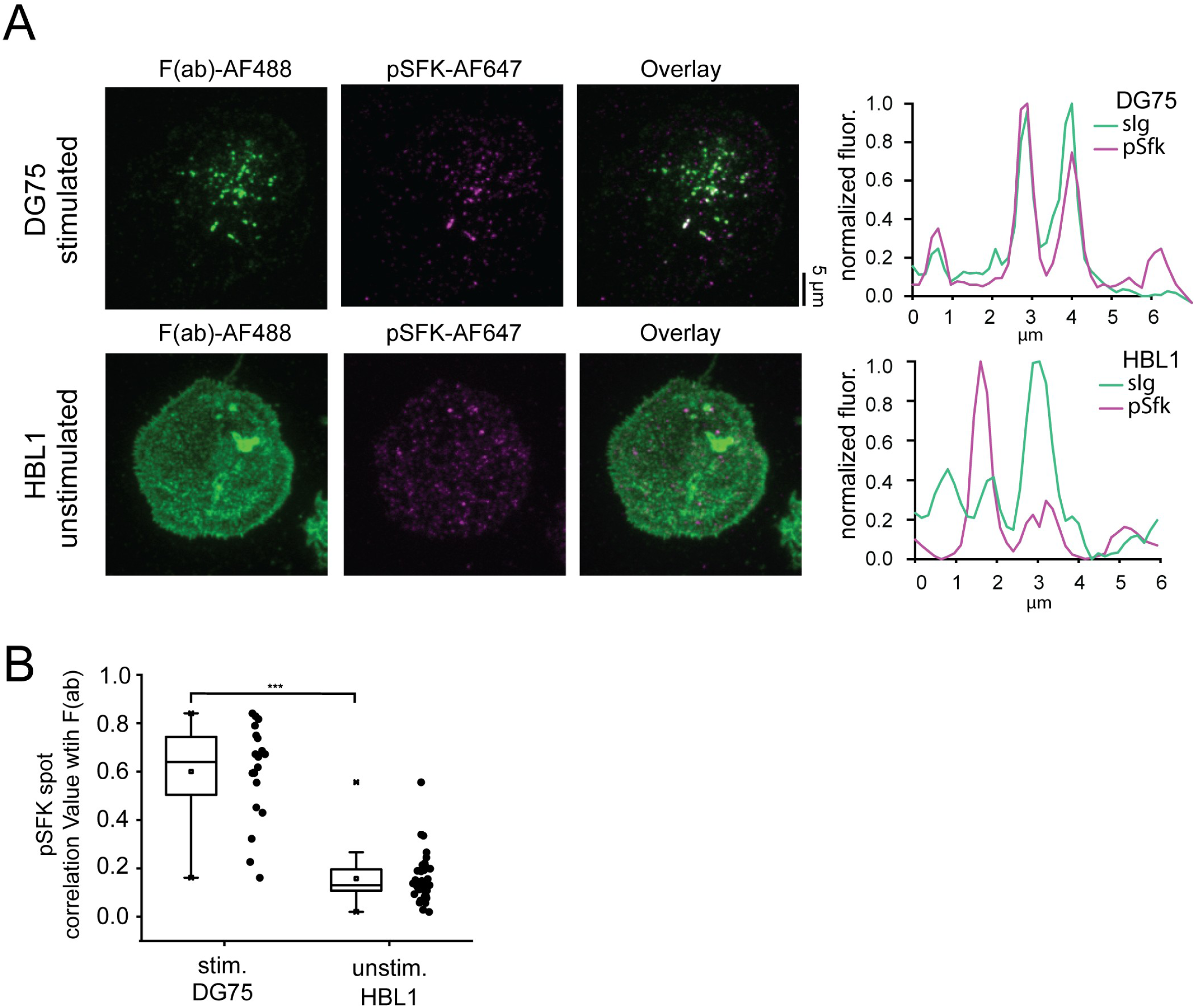
**(A)** TIRF images sIg and pSFK in stimulated DG75 cells and unstimulated HBL1 cells. **(B)** Colocalization values for pSFK spots and sIg. pSFK spots were the reference to identify colocalization with sIg. (For HBL1 cells, n = 40 cells and 2152 spots analyzed from 2 biological replicates. For DG75 cells, n = 20 cells and 326 spots analyzed.)

To investigate the molecular mechanisms of pCD79A disengagement from the BCR complex, we tested whether the AP2-binding deficient CD79B^Y196F^ mutation identified in HBL1 cells might alter localization of CD79A after phosphorylation. CD79A and CD79B are thought to be linked co-receptor proteins that are translated and trafficked to the plasma membrane together, and we hypothesized that this mutation in CD79B might alter the localization of CD79A (33, 34). To test this, we overexpressed GFP-tagged CD79B^Y196F^ in DG75 cells and performed colocalization analysis to determine how well mutant CD79B^Y196F^ associates with the pCD79A after stimulation (Fig. 7 A shows example images). We also overexpressed wild type CD79B (wtCD79B) protein as a control. Previous studies observed that the CD79B^Y196F^ mutant is heterozygous in ABC DLBCL cell lines and patient samples. Thus, for these experiments, we left the wild type CD79B protein expression unperturbed in DG75 cells to more closely mimic the heterozygotic expression pattern of CD79B^Y196F^ found in HBL1 cells (7). DG75 cells expressing CD79B^Y196F^ had a slightly lower correlation with pCD79A, compared to cells expressing the wtCD79B protein (Fig. 7B shows correlation values). This was not statistically significant, and we do not believe the segregation of the pCD79A protein away from the BCR complex is caused by this CD79B^Y196F^ mutation. Thus, it is likely that distinct localization of pCD79A in HBL1 cells is linked to the constitutive nature of its activation, or other mutations or changes in HBL1 cells. Future work is needed to determine the exact nature of this molecular switch and its role in downstream signaling in ABC DLBCL cells.

**Figure 7.**
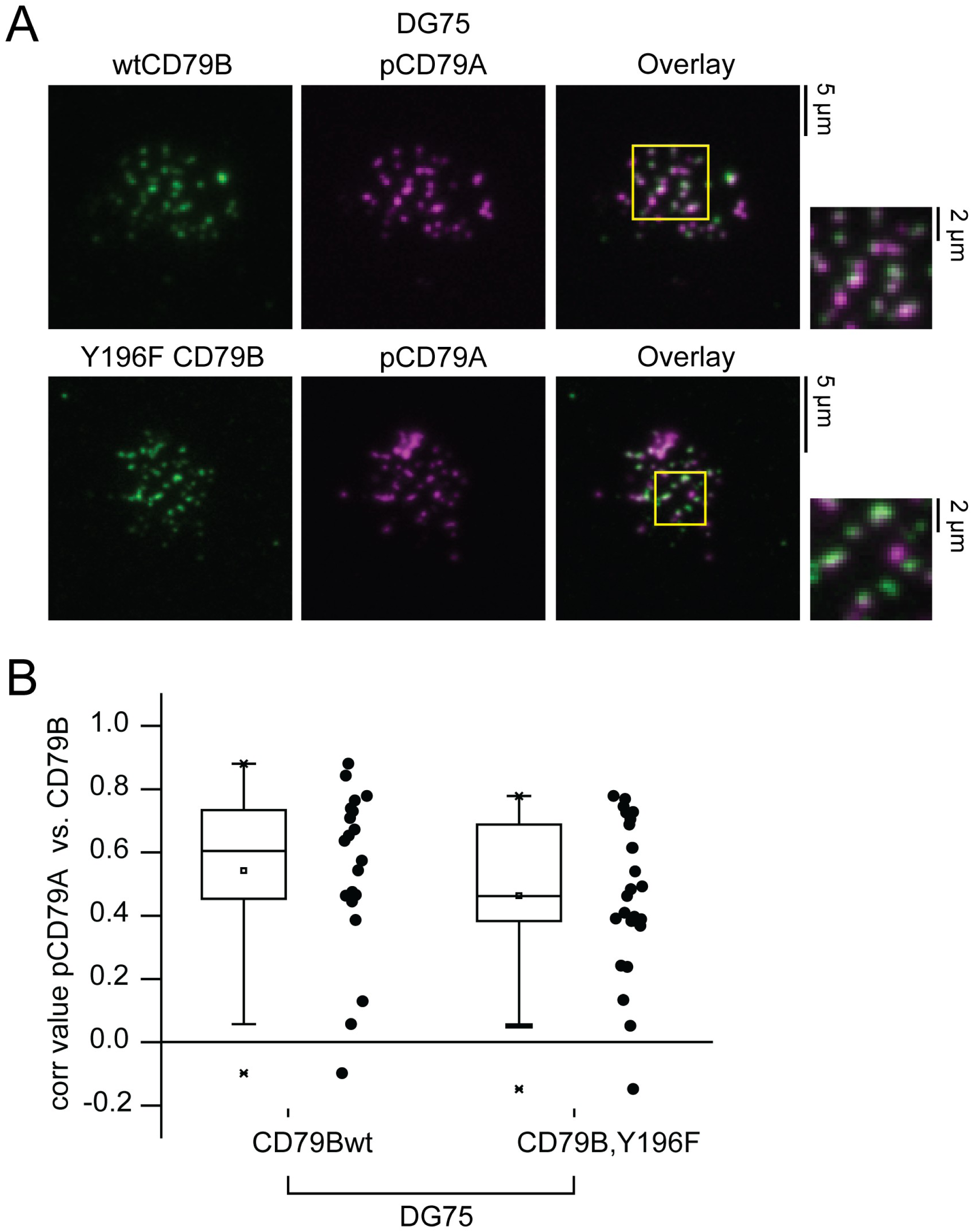
**(A)** TIRF images of DG75 cells overexpressing wild type or mutant CD79B fused to GFP. **(B)** Correlation analysis for pCD79A and CD79B fluorescence. pCD79A spots were the reference to identify colocalization with CD79B. (For cells expressing wtCD79B n = 20 cells and 282 spots analyzed. For cells expressing mutant CD79B n = 25 cells and 354 spots analyzed.)

## DISCUSSION

Our data provide insights into structural mechanisms that underlie the constitutive activation of signaling in ABC DLBCL that contribute to proliferation and cancer. Correlative light and electron microscopy of HBL1 cells revealed that spontaneous clusters of the sIg are localized with cSRMs. cSRM structures are induced in B cells to mediate endocytosis of large activated BCR clusters. Yet, when we imaged pCD79A localization in HBL1s, we observed little colocalization between pCD79A and clathrin lattices on cSRM structures or across the plasma membrane. Based on our data, endocytosis or trafficking of sIg clusters at the plasma membrane are likely affected by improper assembly of activated BCR clusters on cSRMs and their subsequent uptake. Future work to determine whether cSRM structures play a role in maintaining the activated BCR at the plasma membrane is important to determine if drugs that target specific endocytic pathway could be effective treatments for these lymphoma.

Our analysis of activated CD79A at the plasma membrane shows that pCD79A segregates away from the other two components of the trimeric BCR complex (sIg and CD79B) in HBL1 cells. Unphosphorylated CD79A in HLBL1, however, would still be associated in a cluster of slg or CD79B. An alternate interpretation of our data could be that the sIg clusters we observe appear separated from the pCD79A because the F(ab) fragment is staining secreted BCR proteins which have attached to the outer plasma membrane by binding Fcµ Receptors. This is not the case, however, because clustered sIg is highly colocalized with total CD79A and CD79B, indicating that the inactive BCR complex is indeed intact and presented in the plasma membrane. Specifically, secreted BCR proteins contain the sIg (typically IgM) only and would not colocalize with CD79A and CD79B. We therefore propose that CD79A disengages from the membrane embedded BCR complex, possibly triggered by phosphorylation. Importantly, past work has suggested that components of the B cell complex might separate during phosphorylation (35). CD79A/CD79B failed to co-immunoprecipitate with the sIg after antigen stimulation, showing that antigen activation and phosphorylation cause a destabilization of the B cell receptor complex in mouse lymphoma B cells. Our data supports the model that this behavior is common and constitutive in some human ABC cell types. Immune signaling is constitutively active in HBL1 cells, and this complicates identification of the triggering event that leads to pCD79A separation from the sIg and CD79B in ABC cells.

The observation that pCD79A segregates away from the BCR complex in HBL1 cells raises questions about the role of plasma membrane-localized BCR in maintaining pathogenic signaling in lymphoma. First, what is the level of downstream signaling induced by the disengaged pCD79A protein? Our data indicates that pSFK has a similar localization pattern to pCD79A and does not colocalize with the sIg. Thus, pCD79A may act together with other regulators of BCR activation to potentiate downstream activation of NF-κB. Determining other proteins involved in aberrant pCD79A signaling from the plasma membrane may identify novel drug targets. A second question is how widespread this phenotype is among patient samples. Our data show that a second ABC DLBCL cell line TMD8 contains pCD79A that is not colocalized with the sIg. It is, however, important to determine if this phenotype is present in primary patient samples to link this phenotype to the diversity of human diseases found in the clinic.

Finally, the intracellular fate of the separated pCD79A is still unknown. It is possible that pCD79A is eventually endocytosed through clathrin independent mechanisms but the dissociation of pCD79a away from clathrin and slg/CD79B prevents it from trafficking to the correct intracellular compartment. Likewise, pCD79A might maintain signaling even within endosomes, similar to ligand-bound G protein-coupled receptors or Epidermal Growth Factor Receptor (5, 36–38). The fate of these components after they are taken up from the plasma membrane are still unclear and an area for future work (39).

Improving our understanding of how BCR coordinates signaling and endocytosis in lymphoma is key to developing novel interventions for patients with blood cancers. Here, we used nanoscale imaging to characterize interactions between BCR and plasma membrane features in cancerous B cells. These methods may be applied to other cell types to illuminate the effect of specific mutations that are thought to impact endocytosis and their role in the structure of the plasma membrane of B cells.

## METHODS

### Cell Culture and Imaging reagents

DLBCL cell lines (HBL1, TMD8) were obtained from Dr. Louis Staudt’s lab (NCI, NIH). DLBCL cell lines were cultured in RPMI 1640 media without phenol red and with 1% Penicillin / Streptomycin and 10% FBS added. DG75 cells were obtained from ATCC (catalog # CRL-2625). DG75 cells were cultured in RPMI 1640 media without phenol red and with 10% FBS, 1% Penicillin/Streptomycin, 10 mM HEPES pH 7.4, and 1 mM Sodium Pyruvate. Both cell lines were diluted 1:3 every 2-3 days.

A F(ab) fragment linked to AlexaFluor 488 (Jackson ImmunoResearch # 109-547-043) or Alexa Fluor 647 (for STORM imaging; Jackson ImmunoResearch # 109-607-043) was used to stain for the B cell receptor. Cells were spun down at 200xg for 10 minutes and then resuspended at 10×10^6^ cells\mL in media with the F(ab) fragment. Cells were stained on ice for 20 minutes and then washed in cold RPMI media twice before plating on Poly-Lysine coated coverslips. For all imaging experiments, 1.5×10^6^ cells\mL were added to coverslips and allowed to adhere for 10 minutes. Other antibodies used for intracellular staining in imaging experiments are listed below.

|  |  |  |
| --- | --- | --- |
| Phosphor-CD79A (Tyr182) (D1B9) | Cell Signaling Technologies | 29742S |
| CD79A | Cell Signaling Technologies | 3351S |
| CD79B | Invitrogen | PA5-79000 |
| AP2 | Pierce | MA1-064 |
| Clathrin heavy chain (X-22) | Invitrogen | MA1-065 |
| Goat anti-Rabbit IgG, Alexa Fluor 488 | Invitrogen | A11034 |
| F(ab')<sub>2</sub>-Goat anti-Mouse IgG, Alexa Fluor 647 | Invitrogen | A21237 |
| F(ab')<sub>2</sub>-Goat anti-Mouse IgG, Alexa Fluor™ 488 | Invitrogen | A11017 |
| Phospho-Src Family (Tyr416) (D49G4) Rabbit Monoclonal Antibody | Cell Signaling Technologies | 6943 |

### DG75 cell transfection and stimulation

DG75 cells were transfected using the Lonza Nucleofector program M-013 and supplement/solutions from Kit V (Lonza VCA-1003). 5×10^6^ total cells were spun down at 200xg for 10 minutes and then resuspended in Lonza Kit V solutions (82 µl Nucleofector solution + 18µl supplement) with 5 µg plasmid DNA added. After nucleofection 500 µl of warmed media was added to the cuvette and cells were transferred to a T-25 flask with 5mL warmed media. Cells were collected 24 hours after transfection for imaging experiments.

All stimulated DG75 cells were treated using the same protocol. DG75 cells were stimulated with 8 µg/mL F(ab’)2 (Jackson ImmunoResearch # 109-006-127) for 15 minutes at 37°C. Cells were then transferred to ice and spun down at 200xg for 10 minutes before plating on Poly-Lysine coated coverslips for imaging experiments.

DG75 cells treated with actin inhibitors were spun down and resuspended at 1.5×10^6^ cells per 1 mL of RPMI. Cytochalasin D or Latrunculin A was added to cells at a final concentration of 10 or 50 µM for Cytochalasin D and 1µM for Latrunculin A. Cells treated with Cytochalasin D were incubated at 37 °C for 15 minutes and cells treated with Latrunculin A were incubated for 5 minutes. After incubation cells were placed on ice and then plated on Poly-L-Lysine coated coverslips for PREM processing (described below) and imaging using TEM.

### Immunofluorescence Assays

For immunofluorescence staining, cells were fixed in 1% paraformaldehyde (PFA) for 1 minute and then moved to 2% PFA for 20 minutes. Next the sample was washed in PBS and permeabilized in 0.5% Triton x-100 for 2 minutes. Samples were then blocked in 3% BSA for 1 hour before staining with a primary antibody diluted in 1% BSA for 1 hour. Cells were again washed in 3% BSA before staining with a secondary antibody (if the primary antibody was not already conjugated to an Alexa Fluor dye) diluted in 1% BSA for 1 hour. Samples were then washed in 3% BSA and PBS before fixing in 2% PFA for 10 minutes. Finally, cells were washed in PBS before imaging.

### dSTORM

The super-resolution localization method, dSTORM, was used to localize the B cell receptor and AP2 in correlative light and electron microscopy experiments. All dSTORM experiments were done using the Alexa Fluor 647 dye conjugated to a F(ab) fragment, an antibody to the phosphorylated CD79A protein or a secondary antibody. STORM buffer for Alexa Fluor 647 was made fresh for each experiment with 10% glucose in PBS solution. Right before imaging 1.5mL of the glucose solution was mixed with glucose oxidase (final concentration 0.8 mg/mL; CAS 9001-05-2), catalase (final concentration 40 µg/mL; CAS 9001-37-0), and 18.75 µl of 99% β-mercaptoethanol to make the imaging buffer. The imaging buffer was added to the coverslip in an imaging chamber and sealed on top with an empty coverslip. dSTORM imaging was carried out using a Nikon Eclipse Ti2 inverted microscope with a 100x, 1.49 NA objective and an ANDOR iXon Ultra EMCCD camera under the control of Nikon Elements NSTORM software. dSTORM imaging was done in TIRF mode with 647 nm excitation collecting 45,000 frames for the B cell receptor and 30,000 frames for AP-2. dSTORM images were processed using ThunderSTORM and the same settings as previously described (19).

### Super-resolution Correlative Light and Electron Microscopy (CLEM)

Cells were prepped using immunofluorescence or surface stain protocols described above with two modifications for CLEM. First, cells were plated on coverslips that have gold fiducial markers embedded in the glass. Second, cells were unroofed using a syringe to spray stabilization buffer (70 mM KCl, 5mM MgCl_2_, 3mM EGTA, 30 mM HEPES pH 7.4) on the cells after plating on coverslips. dSTORM imaging was performed as described above. After imaging, a 4 mm diameter circle was etched underneath the coverslip where the images were taken and cells were fixed in 2% glutaraldehyde for 20 minutes. Samples were then stained with 0.1% tannic acid for 20 minutes, rinsed in water and stained with 0.1% uranyl acetate. Samples were dehydrated by washing in increasing concentrations of ethanol before critical point drying in a Tousimis instrument (Autosamdri-815). Samples were finally coated in 3 nm of platinum (using rotary coating at 17 degrees) and 5.5nm of carbon in a Leica EM ACE900. Cells imaged in dSTORM were identified in the etched region of the coverslip again before carbon-platinum replicas were lifted from the underlying glass using 5% hydrofluoric acid. Replicas were finally placed on formvar/carbon coated 75-mesh copper grids for imaging on JEOL 1400 TEM using Serial EM software for montaging images. This protocol is roughly the same as described in greater detail in Sochacki et al 2017 (40). dSTORM and TEM images were correlated using PyCLEM (41) and STORM images were transformed to align with scale and orientation of TEM images.

### Analysis of CLEM and TIRF images

TEM images of the inner plasma membrane were analyzed using FIJI to identify clathrin on smooth raised membrane structures. Clathrin structures were identified using PyCLEM for semi-automatic segmentation of EM images (41). Cell maps of cSRM and clathrin structures were generating in FIJI the same as previously described (19). Transformed dSTORM images from CLEM experiments were overlayed on structure maps to determine whether fluorescence overlaps with endocytic structures and to calculate the size of overlapping fluorescent regions reported in Figure 1. The percentage of sIg clusters correlated with cSRMs was calculated using the method described above, except that instead of outputting the size of overlapping regions, the number of overlapping regions was counted and divided by the total number of sIg clusters observed in TIRF images. BCR clusters in TIRF images from Figure 1B were manually segmented using the Li thresholding method in FIJI. After segmenting clusters, area measurements were also calculated in FIJI.

TIRF images in Figures 2, 4, and 5 were analyzed using correlation software written in MATLAB (32). Images taken in the red channel (pCD79A stain) were used as a reference to select points and then determine if fluorescence was present in the green channel (clathrin, sIg or CD79B) to calculate correlation values using Pearson’s correlation coefficients as described previously. The region size for each correlation value was 25 nm. Therefore, the image correlation value at these locations is a measure of colocalized clustering. For whole cell (whole image) analysis the entire cell membrane region was used for Pearson’s correlation measurements.

### Flow cytometry-based sIg uptake assay

BCR internalization assays were performed using the same protocol as previously described (19). Briefly, the extracellular domain of the sIg was stained with a F(ab) fragment attached to FITC or Alexa fluor 647 at 4°C. Cells were then moved to 37°C to allow sIg uptake to continue for up to 1 hour. Cells were then fixed in 2% PFA for 20 minutes and analyzed using flow cytometry. For each time point, 10,000 cells in the live cell gate were collected. As the receptor-F(ab) complex is endocytosed and moved into lower pH endosomes, the FITC fluorescence is quenched. The relative decline in FITC fluorescence corrected for changes in fluorescence that result from dissociation of F(ab) over time (change in Alexa Fluor 647 conjugated F(ab) was measured for this correction) is reported. sIg uptake is calculated using the formula below:

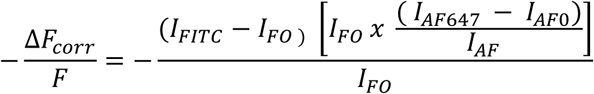

I_FITC_ is the mean fluorescence intensity (MFI) in the FITC channel at the time of measurement, I_F0_ is the initial MFI in the FITC channel, I_AF647_ is the MFI in the Alexa Fluor 647 channel at the time of measurement, and I_AF0_ is the initial MFI in the Alexa Fluor 647 channel.

### Live cell imaging and clathrin residency time analysis

Cells were transfected using the same protocol described above, with a plasmid containing genes for overexpression of the clathrin light chain A protein fused to GFP (Addgene #196912). For live imaging analysis, cells were resuspended in 1mL imaging buffer (130 mM NaCl, 2.8 mM KCl, 5 mM CaCl_2_, 1 mM MgCl_2_, 10 mM HEPES, 10 mM glucose, titrated to pH 7.4 with NaOH) + 0.5% FBS and plated on Poly-L-lysine coated coverslips. Imaging chambers were then set on a stage warmed to 37°C. Images were collected in TIRF mode on a Nikon Eclipse Ti2 inverted microscope with a 100x, 1.49 NA objective, at 1 second intervals over 10 minutes (601 loops).

Image analysis to determine clathrin residency time was done using TrackMate in FIJI. Filtering settings for all video images analyzed were as follows:

- LoG (Laplacian of Gaussian) detector applied
- Only regions with a mean spot intensity greater than 1.5 times the background intensity were analyzed
- A simple lap tracker was selected with link distance parameter set to 0.3
- Filters on tracks: only tracks with a track displacement < 0.33 um and track duration > 20seconds were included in analysis

### Structured Illumination TIRF Microscopy

TIRF SIM images were collected on a GE OMX SR microscope with a 60× 1.42 NA oil immersion objective. DG75 cells were stimulated as previously described and then fixed in 4% paraformaldehyde for 15 min. For staining with pCD79A, the cells were additionally unroofed before staining and then blocked 3% BSA for 1 hour before staining. HBL1 cells were not stimulated before unroofing and staining.

### Statistical analysis

All quantitative measurements were compared using a two-sample T test. P values less than 0.05 were considered significant.

## ACKNOWLEDGEMENTS

We would like to acknowledge Dr. Sebastian Scheich for his generous support providing protocols and materials for imaging and culturing DLBCL cell lines. This research and JWT was supported by the Intramural Research Program of the National Heart Lung and Blood Institute, National Institutes of Health (NIH). ADR is supported by K99GM152952 and R00GM152952 (NIGMS).The contributions of the authors are considered works of the United States Government. The findings and conclusions presented in this paper are those of the authors and do not necessarily reflect the views of the NIH or the U.S. Department of Health and Human Services.

## AUTHOR CONTRIBUTIONS

ADR designed and performed experiments, data analysis and wrote the manuscript. KAS helped with project design, writing software, project development, data analysis, and editing the manuscript. LMS Guided project development, provided DLBCL cell lines and plasmids. JWT oversaw the project, helped with writing software, data analysis, writing, and editing the manuscript.

---

**Supplemental Figure 1.** sIg uptake from the plasma membrane measured using flow cytometry at 10 minute intervals for up to an hour of incubation at 37 °C. (n = 3 biological replicates for HBL1 cells and n = 5 biological replicates for DG75 cells.)

The negative uptake calculated after the 30 minute time point is similar to what is observed when you do the assay with two dyes that do not quench at low pH. And so, after 30 minutes, sIg uptake is undetectable in HBL1 cells.

**Supplemental Figure 2.** Analysis of cSRM density in HBL1 cells treated with drugs to block actin filament formation (Cytochalasin D or Latrunculin A) before EM processing and imaging with TEM.

**Supplemental Figure 3.** Examples of pCD79A and sIg colocalization in untreated HBL1 cells. Yellow boxes highlight regions of the cell where we observed sIg clustering without pCD79A colocalization.

**Supplemental Figure 4.** Colocalization analysis of SIM data. Whole cell colocalization values of the sIg compared to pCD79A in stimulated DG75 or unstimulated HBL1 cells.

