## Supplementary figures and images for "Clathrin recruitment and assembly of phosphorylated B cell receptor is impaired in human diffuse large B cell lymphoma"

### Supplemental Figure 1

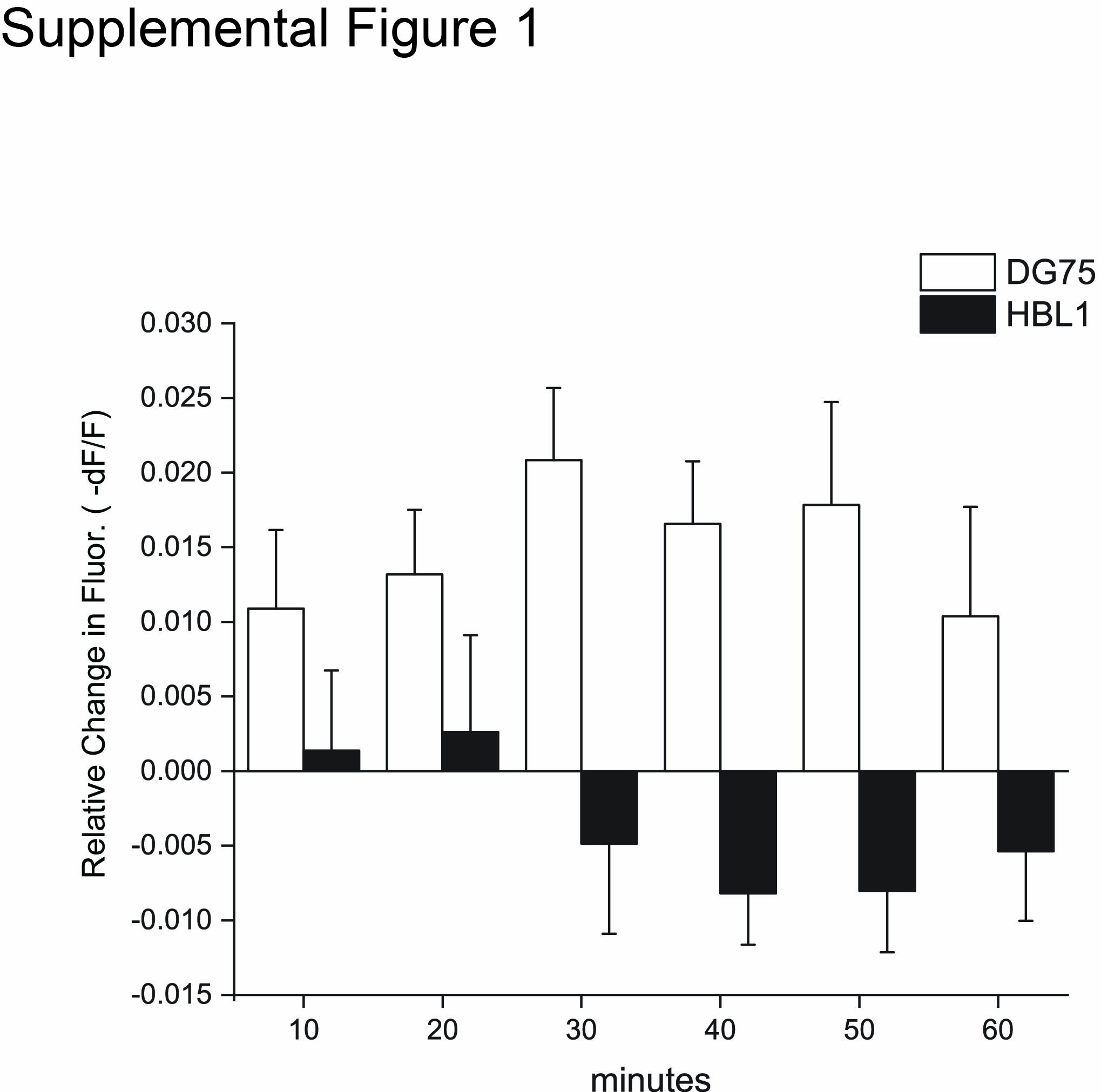

### Supplemental Figure 2

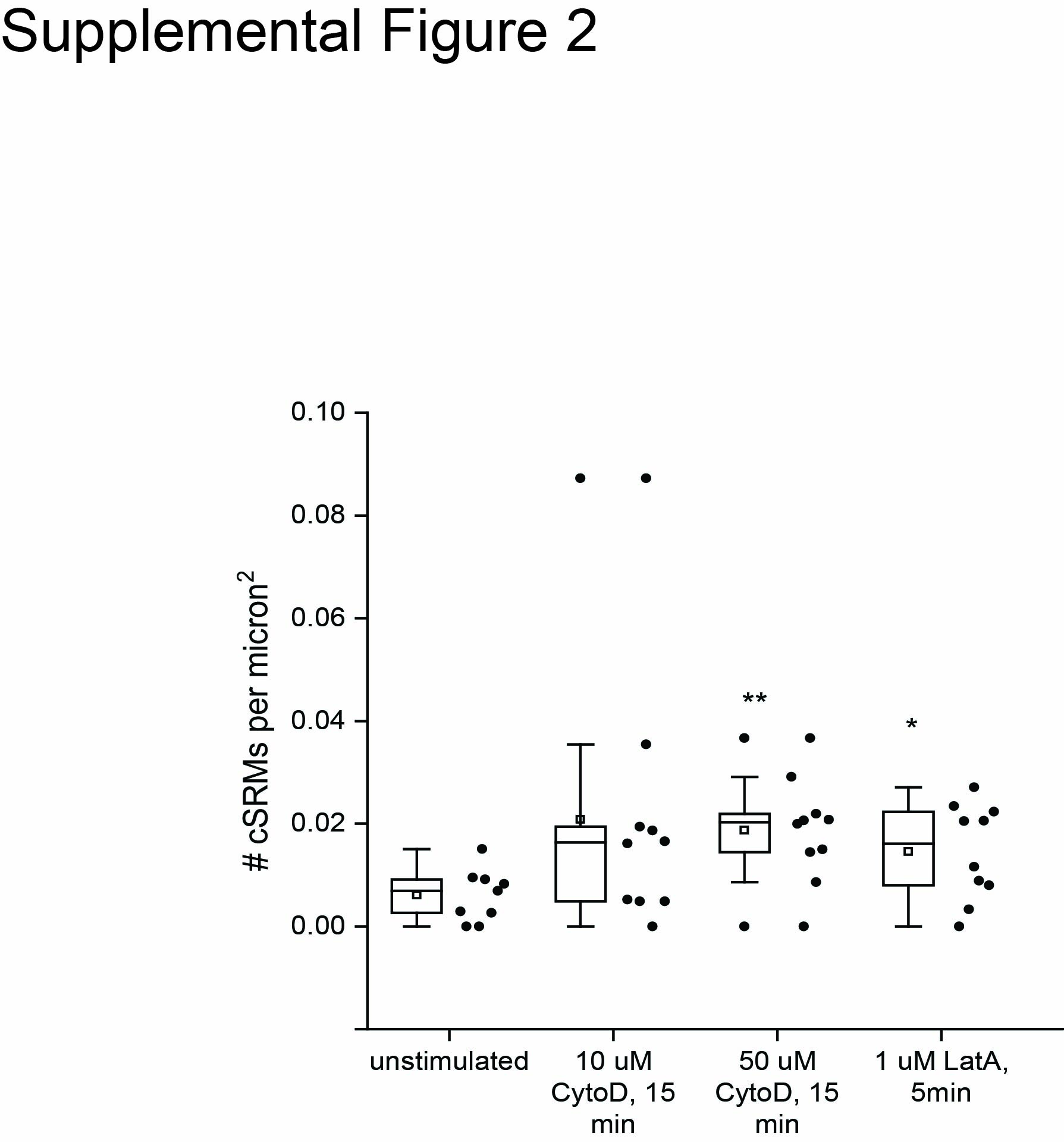

### Supplemental Figure 3

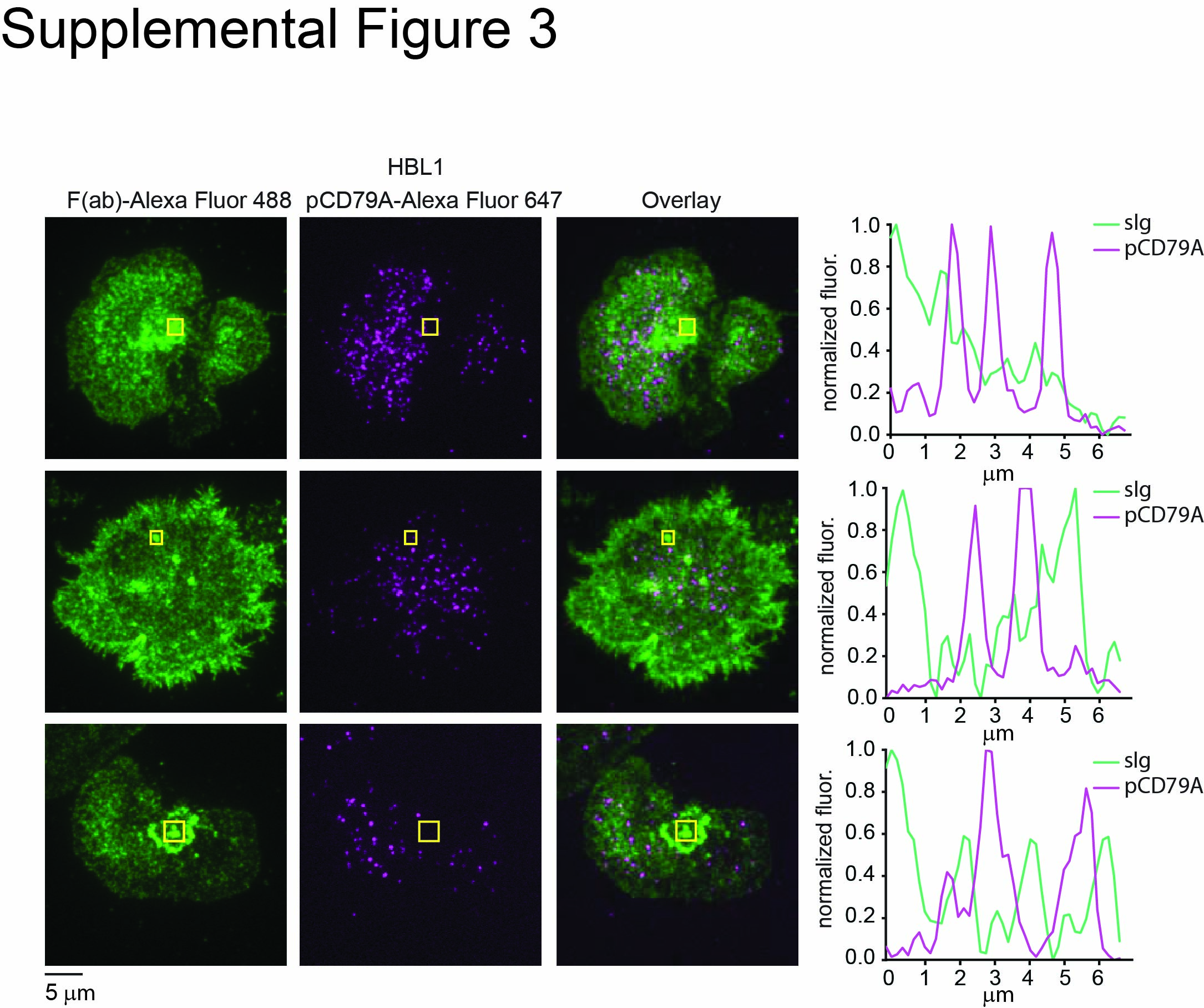

### Supplemental Figure 4

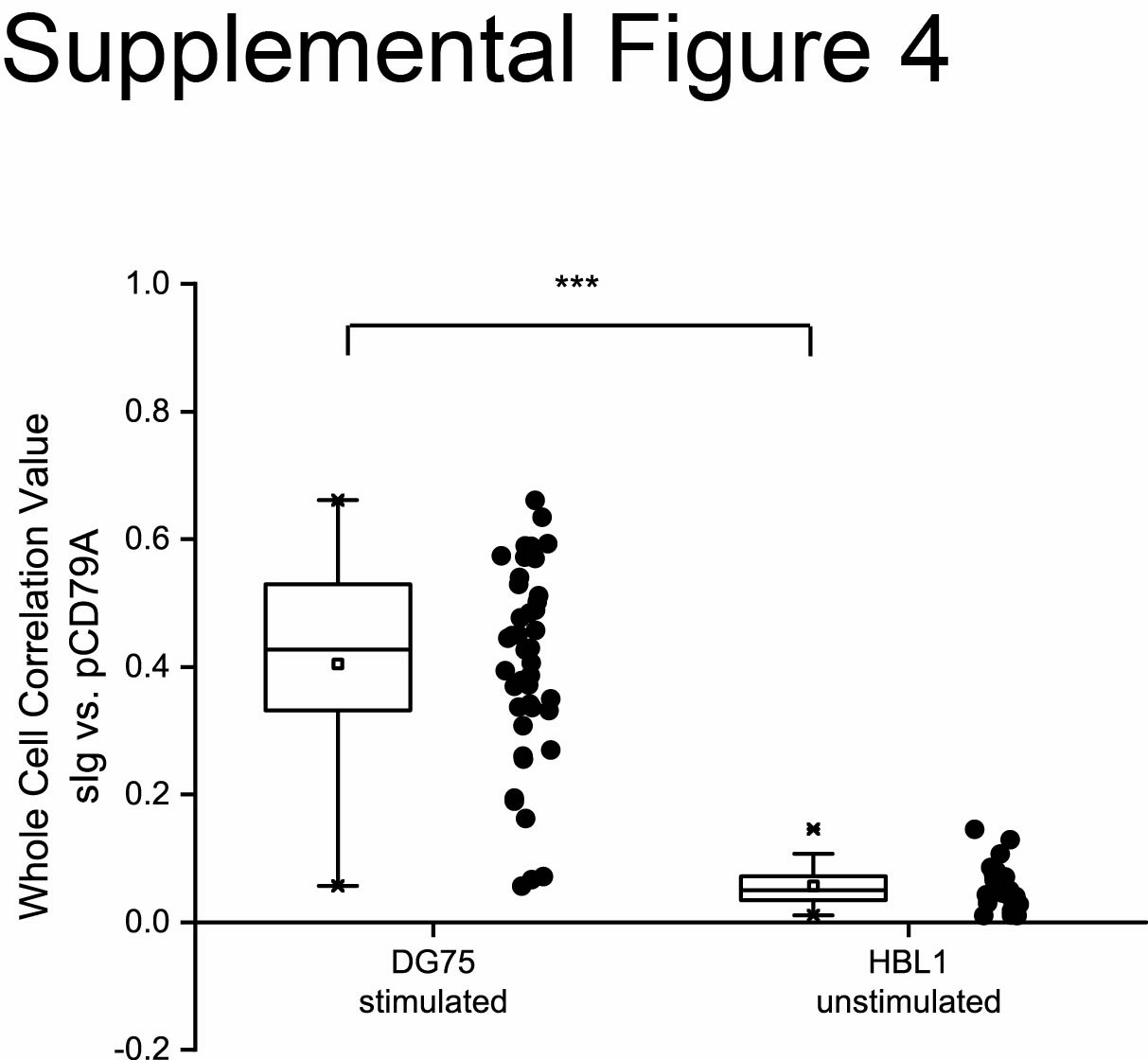
